# RealtimeDecoder: A fast software module for online clusterless decoding

**DOI:** 10.1101/2024.05.03.592417

**Authors:** Joshua P. Chu, Michael E. Coulter, Eric L. Denovellis, Trevor T. K. Nguyen, Daniel F. Liu, Xinyi Deng, Uri T. Eden, Caleb T. Kemere, Loren M. Frank

## Abstract

Decoding algorithms provide a powerful tool for understanding the firing patterns that underlie cognitive processes such as motor control, learning, and recall. When implemented in the context of a real-time system, decoders also make it possible to deliver feedback based on the representational content of ongoing neural activity. That in turn allows experimenters to test hypotheses about the role of that content in driving downstream activity patterns and behaviors. While multiple real-time systems have been developed, they are typically implemented in C++ and are locked to a specific data acquisition system, making them difficult to adapt to new experiments.

Here we present a Python software system that implements online clusterless decoding using state space models in a manner independent of data acquisition systems. The parallelized system processes neural data with temporal resolution of 6 ms and median computational latency <50 ms for medium- to large-scale (32+ tetrodes) rodent hippocampus recordings without the need for spike sorting. It also executes auxiliary functions such as detecting sharp wave ripples from local field potential (LFP) data. Performance is similar to state-of-the-art solutions which use compiled programming languages. We demonstrate this system use in a rat behavior experiment in which the decoder allowed closed loop neurofeedback based on decoded hippocampal spatial representations . This system provides a powerful and easy-to-modify tool for real-time feedback experiments.

## Introduction

The brain enables animals to keep track of information about internal states and the external world and to use that information to guide action selection. This tracking engages neural representations, and thus understanding how those representation relate to internal or external variables can help us understand mental processes.(Knierim 2014). Decoding analyses provide one approach to understanding neural representations: an initial encoding model is built that relates observed variables to spiking, and then this model is inverted to enable predictions of observed variables based on spiking data (Brown et al. 1998). This approach has been used to characterize representations of neural activity from brain regionssuch as the hippocampus (Davidson, Kloosterman, and Wilson 2009; Karlsson and Frank 2009; Pfeiffer and Foster 2013).

The classic application of decoding was to assess how well a given variable could be read out from ongoing neural population activity when that variable (e.g. the position of a limb or of the animal in space) could be observed. When such a correspondence has been established, decoding can also provide insight into representations expressed in the absence of an observable variable. In the hippocampus, for example, the spiking of “place cells” can be decoded both during movement and during periods of immobility. Strikingly, there are times when this spiking corresponds not to current location but instead to other places in the animals environment or even other environments (Carr, Jadhav, and Frank 2011; Foster 2017; Ólafsdóttir, Bush, and Barry 2018; Pfeiffer 2020). Similarly, decoding has also enabled the development of neurofeedback systems, such as brain-machine interfaces, that can translate neural activity patterns into useful outputs (e.g. moving a cursor on a screen or generating speech in patients with paralysis) (Daly and Wolpaw 2008; Luo, Rabbani, and Crone 2023).

Historically decoding hippocampal spatial activity patterns used a decoder which relied on sorted spikes (spikes that can be “clustered” and thereby assigned with reasonable confidence to a single neuron)(Davidson, Kloosterman, and Wilson 2009; Davoudi and Foster 2019; Diba and Buzsáki 2007; Farooq et al. 2019; Grosmark and Buzsáki 2016; Gupta et al. 2010; Karlsson and Frank 2009; Pfeiffer and Foster 2013; Shin, Tang, and Jadhav 2019; Wu et al. 2017; Zheng et al. 2021). In this decoder a Poisson model was used to describe the neural dynamics of individual place cells, where the Poisson rate is directly related to the place field (Brown et al. 1998; Meer, Carey, and Tanaka 2017; K. Zhang et al. 1998).

One disadvantage of the sorted spikes decoder is that it excludes lower amplitude or otherwise non-clusterable spikes. These spikes nevertheless contain valuable information for decoding, and alternative models known as clusterless decoders have been developed. (Deng, Liu, Karlsson, et al. 2016; Deng, Liu, Kay, et al. 2015; Denovellis et al. 2021; Kloosterman et al. 2014; Williams et al. 2020). These decoders use many more of the recorded spikes (typically all that pass a specific amplitude threshold), and provide more accurate decoding compared to sorted spike decoders in cases that have been tested (Deng, Liu, Kay, et al. 2015; Kloosterman et al. 2014). Subsequent studies have applied these methods to derive scientific conclusions (Gillespie et al. 2021; Hu et al. 2018; Michon et al. 2019).

Clusterless decoding thus offers a powerful tool for identifying the content of neural representations, and real-time implementations have the potential to enable the use of this tool in closed-loop experiments. Current implementations are tied to a specific data acquisition system and written in a compiled programming language, however, which increases the difficulty of customization for end users without more advanced programming experience (Ciliberti, Michon, and Kloosterman 2018). We therefore developed `RealtimeDecoder` our software program that implements the state space models in (Denovellis et al. 2021) for online, real-time clusterless decoding (https://github.com/LorenFrankLab/realtime_decoder). The system is parallelized and written entirely in Python for ease of use for both users and developers. Despite using an interpreted language, the software achieves computational latencies similar to state-of-the-art solutions. An added benefit is the implementation of a state space model, allowing the experimenter to use both likelihood and posterior for downstream analysis.

In this work we describe the architecture and performance of `RealtimeDecoder`. We focus on the latencies of computations and feedback, which are especially relevant in a real-time context. We also demonstrate the system’s use in a live closed-loop experiment for proof of concept, and we briefly discuss some results. Our hope is that this work will help advance scientific research by enabling other closed-loop experiments that can elucidate the role of hippocampal spatial representations.

## Materials and Methods

### Model formulation

The clusterless decoder used in this work is based on (Deng, Liu, Kay, et al. 2015; Denovellis et al. 2021). Similar to the sorted spikes decoder, this decoder uses a Bayesian framework. The model is governed by the equation

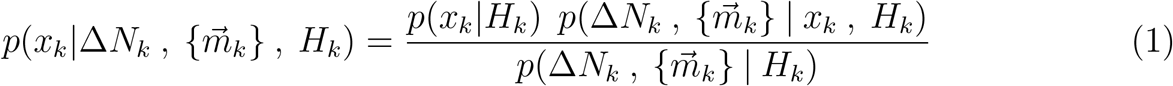

where for a given time bin *k, x*_*k*_ is decoded position, Δ*N*_*k*_ is the number of spikes emitted, and 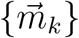 is the set of marks of length Δ*N*_*k*_, that is, vectors associated with each spike observed in time bin *k*. In practice the marks used in the model are spike waveform features such as peak amplitudes. Lastly *H*_*k*_ represents the spiking history from time 1 to time *k*.

The distribution *p*(*x*_*k*_|*H*_*k*_) in Eq. 1 is given by

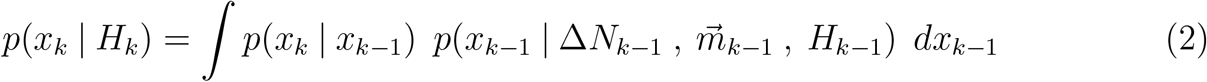

assuming that (1) transitions from *x*_*k−*1_ to *x*_*k*_ are Markovian, and (2) the distribution of the current decoded position is independent of past spiking history, given the previous decoded position (Deng, Liu, Kay, et al. 2015).

For the clusterless likelihood, the relation is

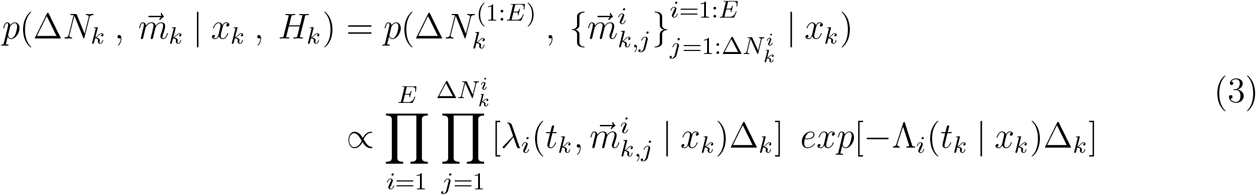

where 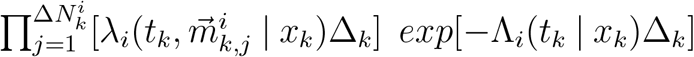 is the likelihood for electrode group *i* = 1, …, *E*. (Our model assumed 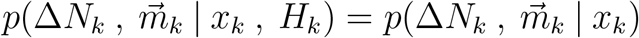. For each electrode group, the product 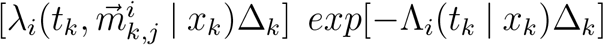 is computed for every spike *j* within time bin *k*. If there are no spikes observed for electrode group *i* in time bin *k*, then that electrode group’s likelihood becomes

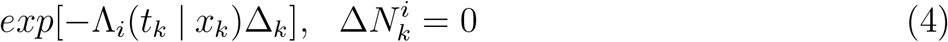

The likelihood consists of intensity functions

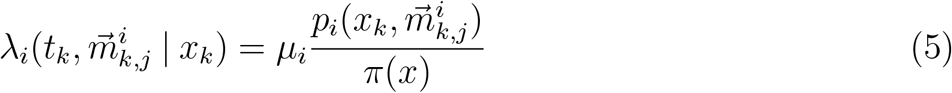

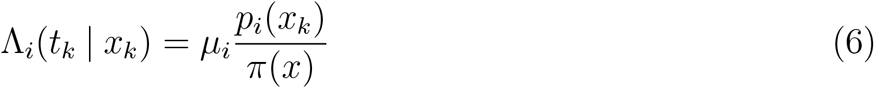

where *µ*_*i*_ is the mean firing rate of electrode group *i* and *π*(*x*) is the occupancy, a distribution of positions the animal has visited. *p*_*i*_(*x*_*k*_) is the probability of observing any spike given for electrode *i*, given the position. Lastly 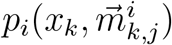 represents a joint probability distribution over position and marks. During the model learning phase, no computation is performed–each mark 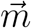 is simply saved along with the position x at which the mark was observed. Once the learning phase is complete, 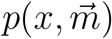 is estimated on-the-fly for each observed mark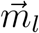. First a weight is computed for each mark 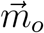 that has been saved in the model, using a Gaussian kernel:

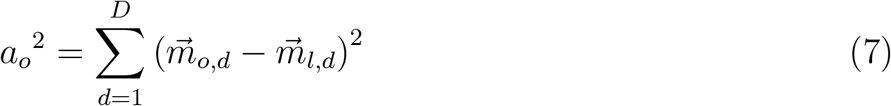

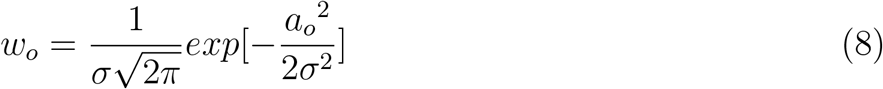

where *D* is the number of mark dimensions and *σ* is a user-defined parameter. Then a weighted histogram representing the estimate 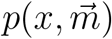 is computed using the positions {*x*_*o*_} saved during the learning phase, and the corresponding weights {*w*_*o*_} described above.

Finally the posterior distribution can be written as

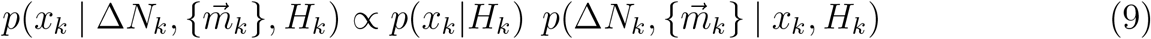

where *p*(*x*_*k*_|*H*_*k*_) is given in Eq. 2 and 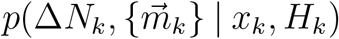 is given in Eq. 3.

### Software architecture

Since the clusterless likelihood in Eq 3 is a product over electrode groups, this formula presents a straightforward scheme for parallelization. Each factor in can be computed simultaneously before being combined into the overall likelihood. The likelihood is then used to estimate the posterior distribution in Eq. 9.

To carry out these computations `RealtimeDecoder` uses three different input data streams: LFP, spike events (particularly the waveform data), and position. An overview of the computational approach is shown in Fig. 1. Input data (ellipses) flow to parallel processes (rounded rectangles) which compute intermediate quantities (rhomboids), resulting in a reward trigger output (parallelogram) if a specific, experimenter specified representation is detected. To send messages between different processes, we used the low-latency messaging protocol MPI (Walker and Dongarra 1996).

**Figure 1.**
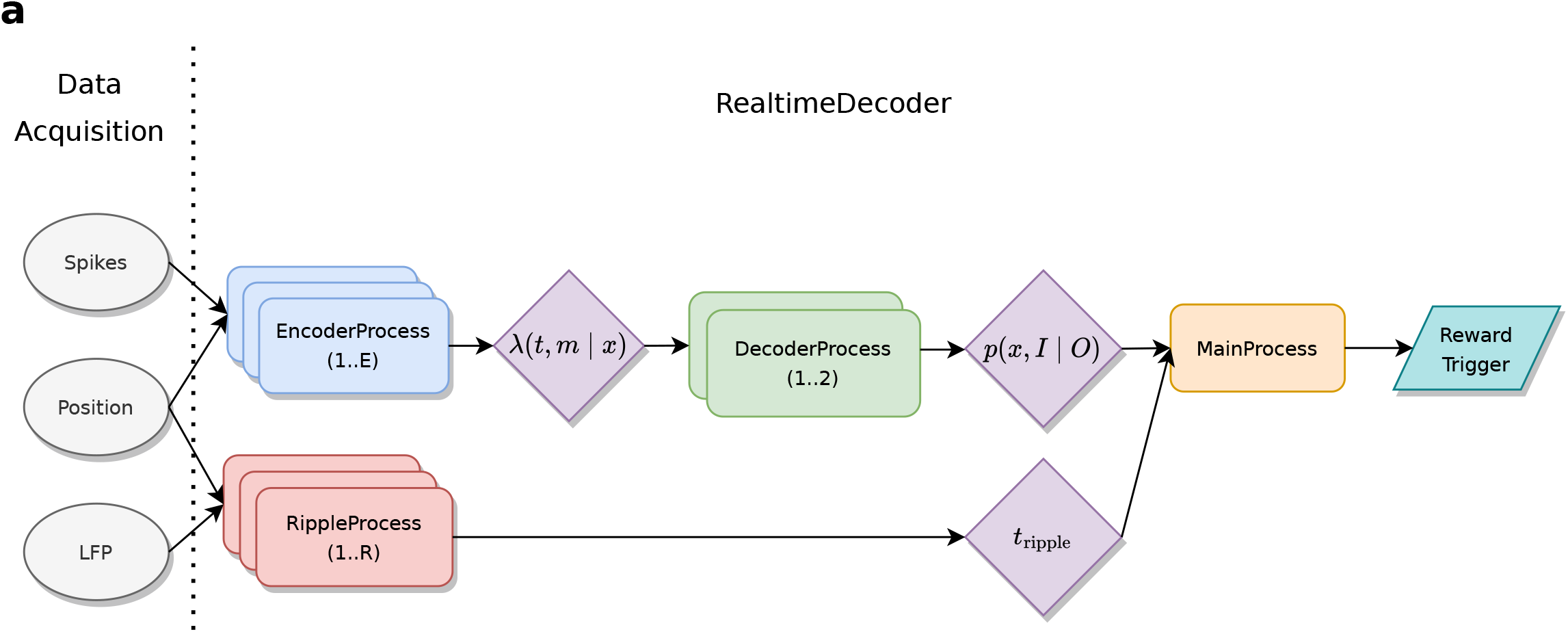
(a) Data flows through different Process nodes in the `RealtimeDecoder` system, which are responsible for different computations.

The primary objective of online real-time clusterless decoding is to estimate the posterior distribution in relation 9, which is a conditional probability distribution over a variable x. In this application x represents linearized (1D) position. Recall that the clusterless likelihood in Eq. 3 is a product over electrode groups, which presents an obvious target over which to parallelize. The overall software architecture reflects this feature, as the computation of Eq. 3 can be split among multiple processes to reduce total system latency. These sub-computations is then be aggregated to form the posterior probability distribution.

Additionally the system can process LFP data to extract time boundaries in which LFP events (specifically sharp wave ripples or SWRs) occur. Experimenters thus have multiple options when using the system; for example they may specify that only replay with a concomitant SWR will trigger neurofeedback.

`RealtimeDecoder` is implemented as different `*Process` objects, namely `MainProcess`, `RippleProcess`, `EncoderProcess`, and `DecoderProcess` which serve different computational functions (Fig 1). Each instance of these objects (with the exception of the event-driven `GuiProcess`) implements a polling while loop which typically consists of the following pseudocode:

#### MainProcess

The `MainProcess` coordinates all other Process instances by gathering information about the binary data they output to file and monitoring their status to check for runtime errors, among other functions. Inside the `MainProcess` runs a custom data handler that processes data computed by other `*Process` objects, such as position information, ripple onset times, a new posterior distribution estimate, etc. In our usage this object detects replay events; upon detection it sends a signal to dispense reward.

**Listing 1.**
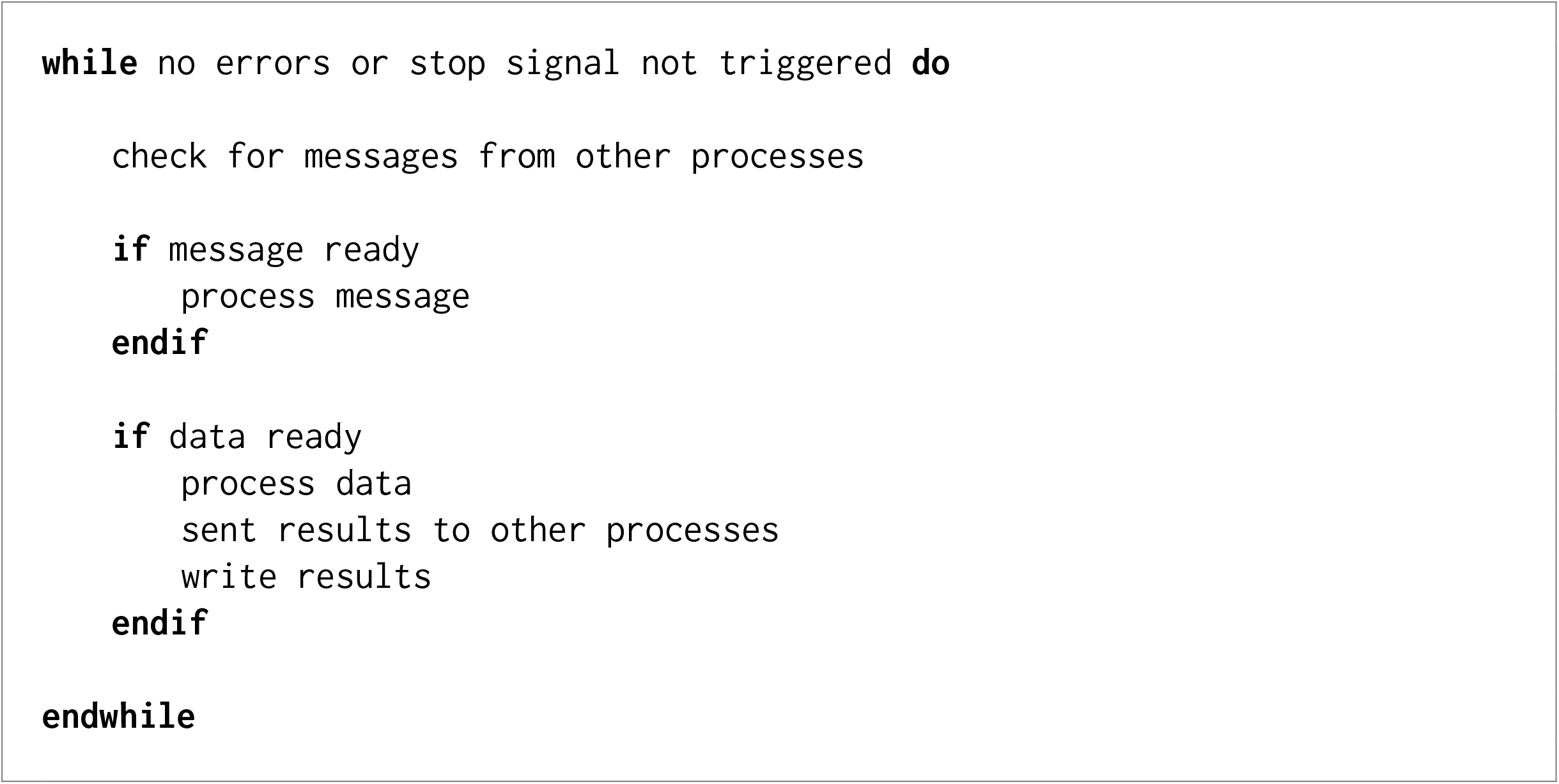
Pseudocode example for a `Process` object

#### RippleProcess

An instance of a `RippleProcess` processes LFP data and is primarily responsible for detecting sharp wave ripples. This occurs via the following procedure: (1) LFP from the data acquisition system is band-pass filtered to the SWR frequency band (150-250 Hz), (2) an estimate of SWR power is obtained by applying a low pass filter to the square of the band-pass filtered data, and finally (3) the baseline (mean) power is estimated. The start of a SWR is then marked when the z-score of this power estimate exceeds a user-defined threshold (typically at least 3).

#### EncoderProcess

An instance of an `EncoderProcess` manages the encoding model for one or more electrode groups (e.g. a single tetrode). Typically if the number of available threads permits, a single instance will handle just one electrode group for minimum computational delay. During the training period, an `EncoderProcess` adds spikes to the encoding model and also computes an estimate of the joint mark intensity function (Deng, Liu, Kay, et al. 2015). When the training period is over, spikes are no longer added to the encoding model, but the joint mark intensity function continues to be estimated.

#### DecoderProcess

The `DecoderProcess` gathers the joint mark intensity functions sent by instances of an `EncoderProcess`. At every user-defined time bin step, it computes the likelihood and posterior distribution of the clusterless decoding state space model. It sends these estimates to the `MainProcess` to be further processed by a custom data handler. This data handler is developed according to the needs of the particular experiment and may implement features such as remote spatial representation detection.

#### GuiProcess

The `GuiProcess` increases user-friendliness of the system. It consists of a visualization window that displays the likelihood, posterior, and state probabilities in real time. It also includes a dialog window that allows the user to change parameters during the course of the experiment, as well as some control options to start and stop the software as a whole.

### System characterization

The utility of an online decoding system for real-time feedback depends in large part on its latency. For example, if one chooses too small of a time step to update the posterior distribution estimate, the system cannot compute the necessary quantities quickly enough to be suitable for real-time use. The overall latency stems from the latencies of four major components (Fig 2a): (1) Network latency, (2) spike incorporation latency, (3) posterior computational latency, and (4) event detection latency, which are further expanded in panel (b). Each of these latencies is expanded in further detail below.

**Figure 2.**
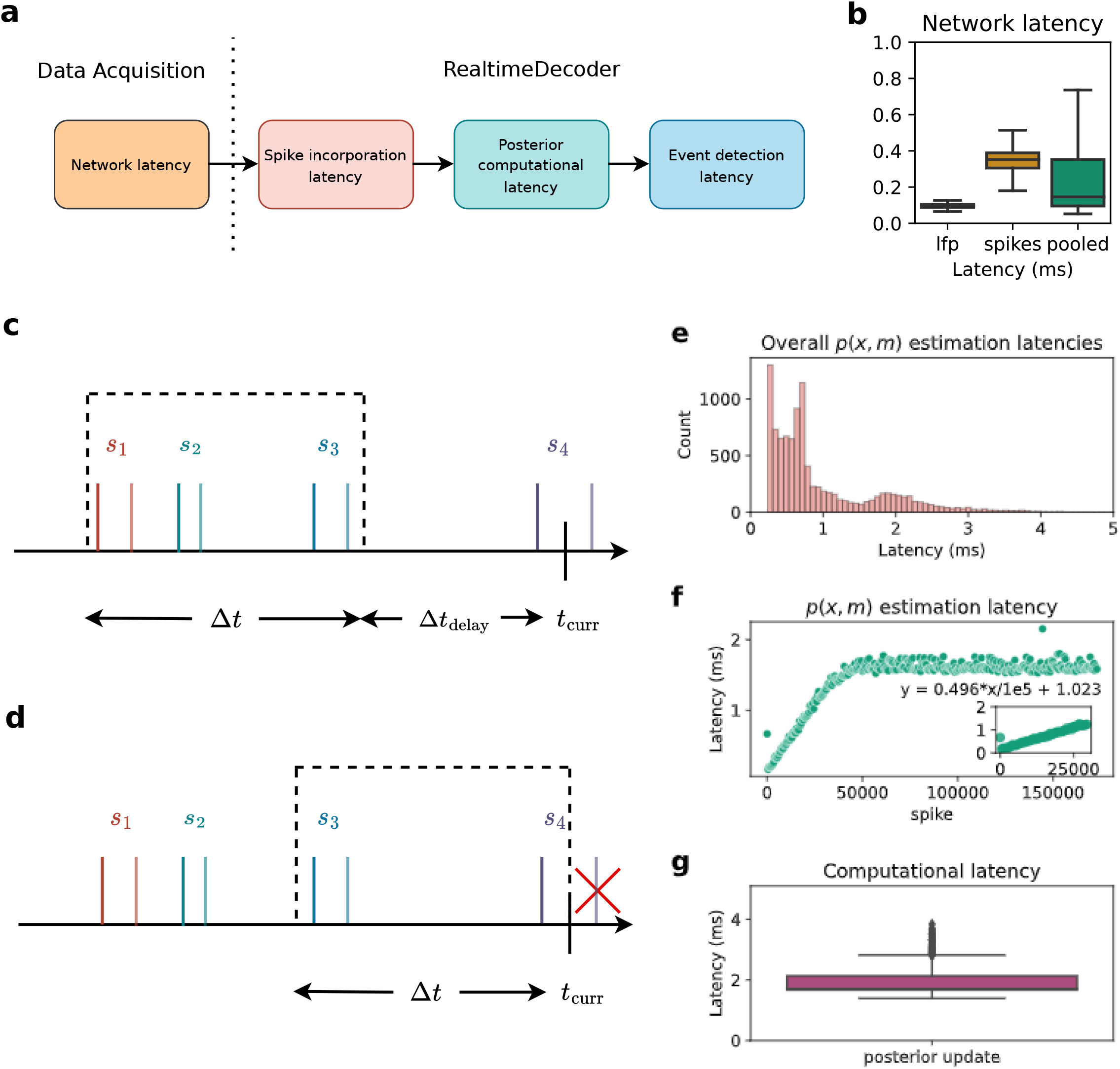
(a) The four major components contributing to overall latency. (b) Network latency. (c)-(d) Four spikes are shown. The darker shade indicates when a spike occur, and the lighter shade indicates when it is in usable form for a `DecoderProcess`. Note that spike 4 is in usable form subsequent to *t*_*curr*_. (c) Time bin for when Δ*t*_*delay*_ > 0. (d) Time bin for when Δ*t*_*delay*_ = 0. (e) Overall distribution of latencies due to estimation of *p*(*x, m*) using an example tetrode. (f) *p*(*x, m*) estimation latencies as a function of number of spikes in the encoding model. (g) Computational latency induced by update of the posterior distribution.

#### Network latency

The network latency (Fig 2b) is the time difference between when data (spikes, LFP, position) is sent out by the data acquisition system, and when that data is actually requested by `RealtimeDecoder`. This type of latency depends on different factors such as network speed, the type of machine running the decoder, etc. Spikes have a higher latency than LFP due to its size, as the entire spike waveform/snippet is transmitted over the network. Nevertheless the median network latency of the pooled data (spikes and LFP combined) is <1 ms when the data acquisition program executes on the same host machine as `RealtimeDecoder`.

#### Spike incorporation latency

The spike incorporation latency refers to the maximum length of time between when a spike occurs and when it is sent to the decoder in a usable form. To be usable the joint mark intensity must be estimated with the spike and be visible to a `DecoderProcess` instance. At that point in time, the spike is ready to be incorporated into the posterior distribution estimation via an update of the likelihood.

The spike incorporation latency is a user-defined parameter and therefore a constant value. It must be long enough to account for the latency incurred from estimating the joint mark intensity function, a step whose computation time typically increases with the number of spikes used to generate the encoding models utilized by the decoder.

In `RealtimeDecoder`, the estimation of the posterior distribution occurs one time bin at a time. The data incorporated into each estimation step are those contained within a certain time window. This window is directly impacted by Δ*t*_*delay*_ and Δt, the time bin size.

*t*_*curr*_ refers to the time at which an update to the posterior occurs. Δ*t*_*delay*_ (a non-negative value) refers to how far behind *t*_*curr*_ should define the upper edge of the window. A combination of Δ*t*_*delay*_ and Δt defines the lower edge. In other words, the window is defined by [*t*_*curr*_ *−* Δ*t*_*delay*_ *−* Δ*t, t*_*curr*_ *−* Δ*t*_*delay*_). The spike incorporation latency is the sum Δ*t*_*delay*_ + Δ*t*.

Fig 2b-c illustrates the effect of the spike incorporation latency on the boundaries of the time window when an update to the posterior is requested at time *t*_*curr*_. In Fig 2c, Δ*t*_*delay*_ is chosen to be a non-zero value so the upper edge of the window lies at a time prior to *t*_*curr*_. Spikes s_1_, s_2_, and s_3_ are incorporated into the update of the posterior.

If Δ*t*_*delay*_ = 0 (Fig 2d), then the upper edge of the window is simply *t*_*curr*_. However, spike s_4_ is NOT incorporated into the posterior update because it has not been transformed into a usable form until a time past *t*_*curr*_. For this reason, it is advised to set Δ*t*_*delay*_ > 0.

Fig 2e-f illustrates some considerations for determining an appropriate value to set as the spike incorporation latency. The joint mark intensity (JMI) in Eq. 5 must be estimated, where estimation of *p*(*x, m*) is the most computationally expensive component. Fig 2e shows the overall distribution of the *p*(*x, m*) estimation latencies. However, this is not the complete picture: Fig. 2f shows that these latencies are a function of number of samples in the encoder model. It thus illustrates typical operation of `RealtimeDecoder`: at some point the encoding model is considered trained, so no more spikes are added to the model and the estimation latency no longer increases linearly. On our test machine the estimation latency increased at an approximate rate of 0.5 ms additional latency for every 10000 additional spikes in the encoding model.

In general, the more spikes are expected to be added to the encoding models, the higher the user must set the value of the spike incorporation latency. Too low of a value would mean some JMI’s cannot be computed and incorporated into the posterior estimation step in time, which could adversely affect the quality and accuracy of the posterior.

#### Posterior computational latency

The posterior computational latency is the computation time required to estimate the posterior distribution for a single update step. Fig 2g shows the distribution of this latency (median value 1.722 ms on our test machine). The latency is affected by multiple factors, including how many electrodes `RealtimeDecoder` is configured to analyze and how many states are in the state space model. The distribution in Fig 2g is useful for informing the user what the time bin size Δ*t* should be. We advise setting Δ*t* to at least the 75th percentile of this distribution. If Δ*t* is too small, then the decoder would not be able to estimate the posterior quickly enough and would fail to operate in real time.

#### Event detection latency

The event detection latency is a user-defined parameter relevant for replay detection. Similar to the spike incorporation latency, it is a constant value. At each time bin step, a window is drawn. If the current timestamp is *t*_*curr*_, then the event detection window is defined by [*t*_*curr*_ *−* Δ*t*_*event*_, *t*_*curr*_] where Δ*t*_*event*_ is the event detection latency, a positive value denoting the size of the window. For example if Δ*t*_*event*_ is 20 ms, then a putative representation will have been expressed for at least 20 ms before it is detected by the software. A higher value of the event detection latency will theoretically reduce the number of spurious detections, at the cost of increasing the reaction time to a true event.

#### Summary

Overall our decoder performs comparably to state-of-the-art solutions (Table 1) despite being written entirely in Python.

**Table 1.** RealtimeDecoder latency performance relative to the decoder in (Ciliberti, Michon, and Kloosterman 2018).

| Decoder | Network latency | Spike incorporation latency | Posterior computation latency | Event detection latency |
| --- | --- | --- | --- | --- |
| Ciliberti 2018 (32 tetrodes) | 0 ms (median, localhost) | 10 ms | ~2.2 ms (median) | 30 ms |
| This work (49 tetrodes) | 0.102 ms (median, localhost) | 12 ms | 1.722 ms (median) | 30 ms |

### Scalability

The estimation of the JMI depends on the mark dimensionality, where Fig. 3(a) demonstrates this relation. Here test inputs were generated deterministically according to Listing 2, where all spikes were identical in magnitude and the distribution of positions was uniform. Note that different test inputs are not expected to change the relations demonstrated by Fig 3(a) since the actual values of the marks and position do not matter when only latency measurements are of concern.

**Listing 2.**
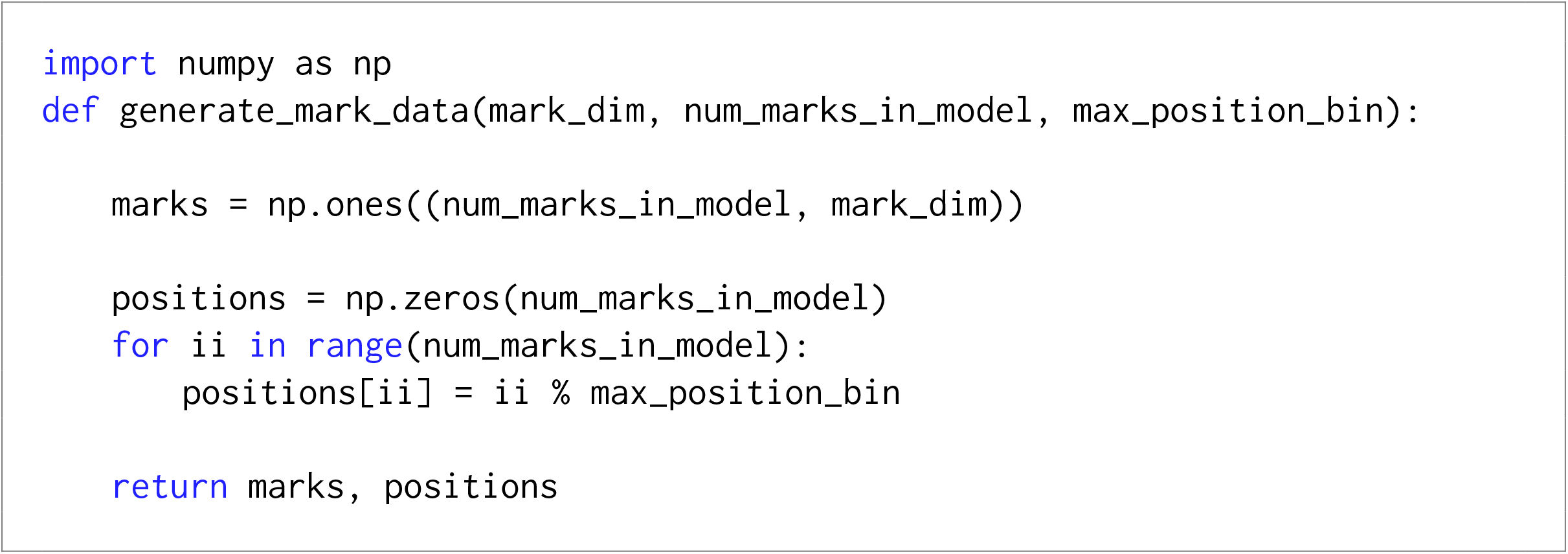
Test input generation for characterizing effect of mark dimensionality on computational latency.

**Figure 3.**
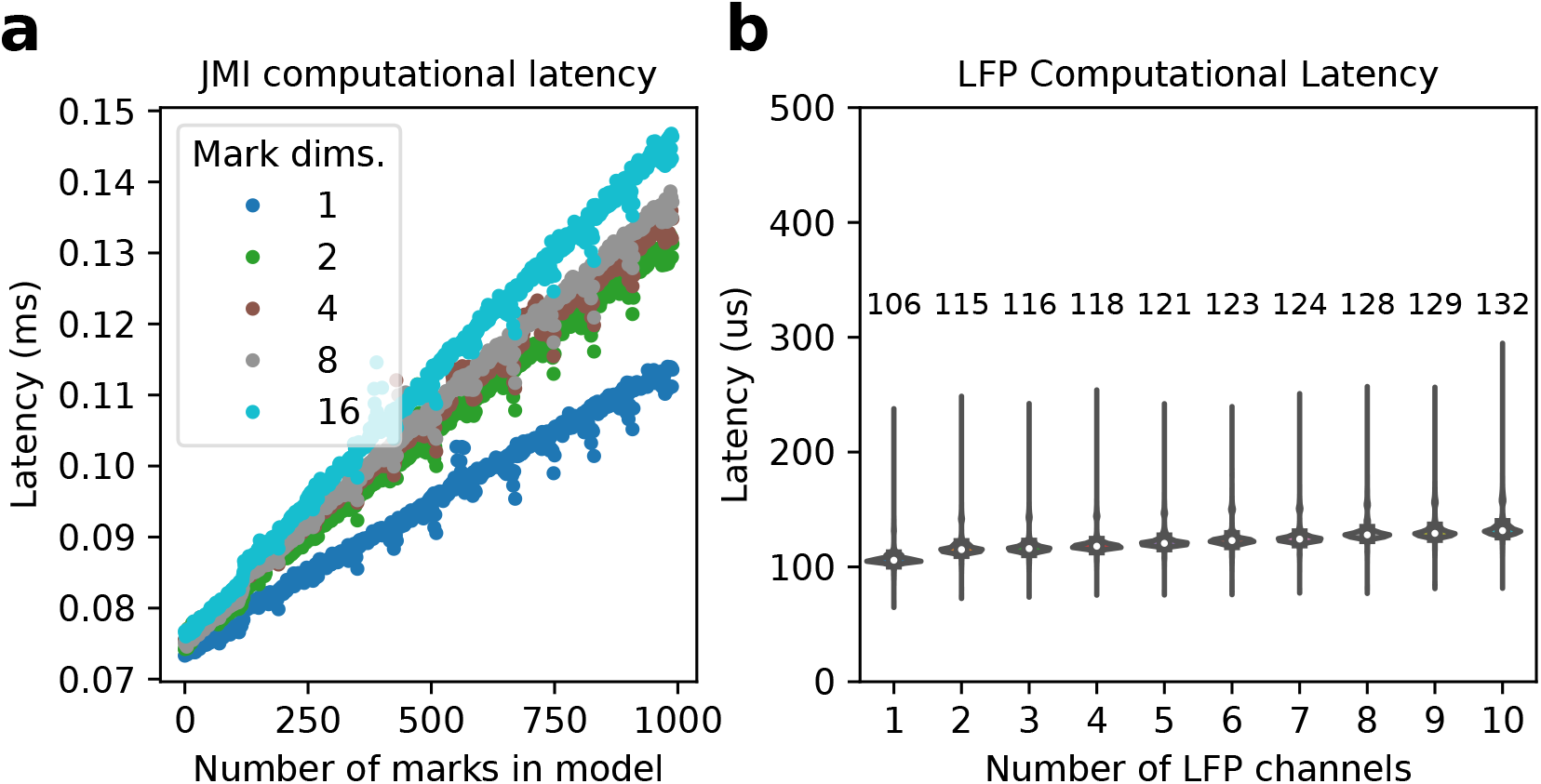
(a) Computational latency for estimation of the JMI depends on the mark dimensionality. (b) LFP computational latency. Median values are labeled above the distribution they were computed from.

Although the computational latencies increase with the number of mark dimensions, the multiplier for this increase is less than the ratio of a given mark dimensionality to a reference mark dimensionality. As an example, increasing the dimensionality from 1 to 16 does not result in a 16X increase in the computational latencies. This is a favorable characteristic especially for experimenters using larger electrode groupings, such as for polymer or silicon multielectrode arrays.

For the LFP processing path, users may want to process multiple channels on one given `RippleProcess` instance. Real-time constraints dictate that every LFP sample must be processed at Δ*t*_*LFP*_ < 1/*fs*_*LFP*_ where Δ*t*_*LFP*_ is the LFP processing time and *fs*_*LFP*_ is the LFP sampling rate.

A `RippleProcess` instance is responsible for processing LFP by filtering the data into the ripple band, estimating the ripple power (or a proxy of it), and determining the start and end times of a ripple. Fig 3(b) shows the computational latency caused by processing LFP data, for a single `RippleProcess` instance. Here the LFP sampling rate was 1500 Hz, so the maximum LFP processing latency is 667 microseconds. These results demonstrate that 10 (and likely more) LFP channels may be processed with comfortable margin to meet the 667 microsecond real-time deadline. Recall that each `*Process` instance, with the exception of `GuiProcess`, implements a polling `while` loop so this deadline represents the upper bound since other tasks must be executed for every loop iteration.

Overall Fig 3 demonstrates favorable scaling characteristics for higher dimensionality data.

## Results

To demonstrate the utility of `RealtimeDecoder` we applied it to an experiment to study internally driven memory retrieval. Specifically, we used `RealtimeDecoder` to deliver reward based on decoded spatial representation to test whether rats can control the activation of specific remote spatial representations . This endeavor is a type of experiment that can decode neural activity while it is occurring and react on a timescale of 10s of milliseconds, bringing researchers to a closer understanding of memory-related neural firing patterns.

In this experiment a Long-Evans rat was surgically implanted with a microdrive with 64 individually adjustable tetrodes (all animal surgery and care was performed in accordance with UCSF IACUC guidelines). Over the course of several weeks, tetrodes were lowered into the CA1 region of dorsal hippocampus. The rat was food-restricted and trained to run on a two-arm maze where it collected sweetened milk reward from lick ports in the maze.

Once tetrodes were in place, the experiment began. Each day consisted of interleaved RUN and REST sessions in the following manner: REST 1, RUN 1, REST 2, RUN 2, REST 3, RUN 3, REST 4. During REST sessions, the animal was placed in an enclosed box away from the maze.

The structure of a RUN session was sub-divided into two tasks (Fig 4a), but the decoder itself runs for the entire duration of a session. During task 1 (∼ 10 min.), a light cue directed the rat to explore each arm (12 visits each). The data collected from this exploration period, and specifically spikes that occurred during periods of running, defined the encoding models contained in the decoder.

**Figure 4.**
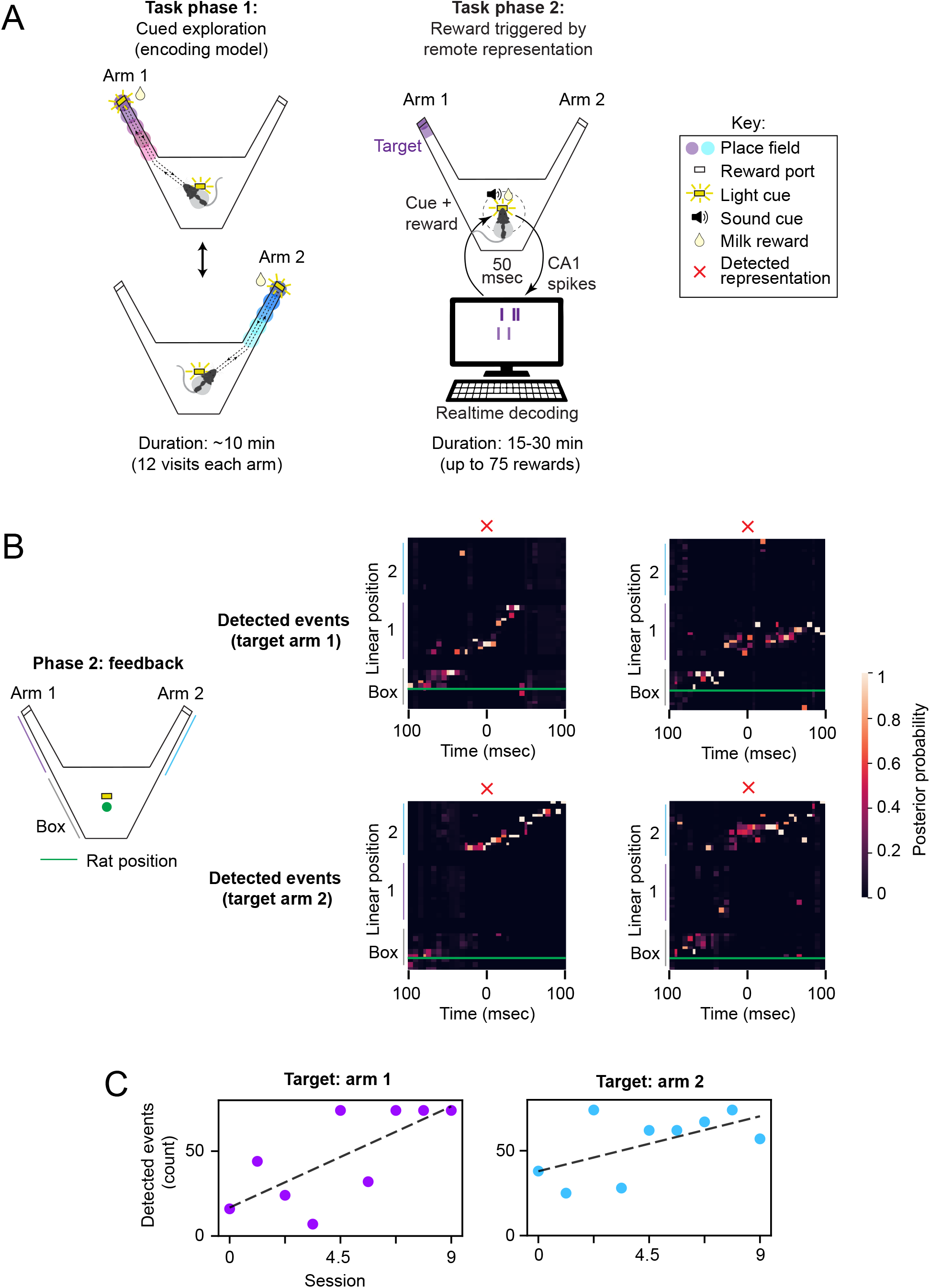
(a) Overview of experiment to which the real-time decoder was applied. Each session consists of two tasks. In task 1 an animal explores the maze arms. This data is used to train the encoding models. In task 2 the animal receives reward for generating a remote representation event. Although place cells and remote sequences are expected to be present, they are not used directly by the clusterless decoder. (b) Example events captured by the decoder. The blue trace is the actual position of the animal. A 100 ms window is drawn around the time of event detection. (c) Number of rewards for each session of neurofeedback.

In task 2, the animal no longer collected reward by running to the end of a maze arm. Instead, reward was only available at a reward port at the center of the maze, located away from each arm. When the decoder detected a remote representation of a target location unknown to the rat (the end of one arm), a sound cue was played and the rat had 3 seconds to nose-poke and collect the milk reward. Importantly, remote representations were only rewarded if the rat was near the center port, away from the arms. Detected representations were required to be high-fidelity events i.e. with little representation in the off-target arm.

To detect a remote representation, a sliding window is applied to the posterior or likelihood, which are both normalized to be a probability distribution. The distribution used for detection is a user-defined parameter. Whenever the distribution is updated (i.e. one time bin step has advanced), a window [*t*_*curr*_ *−* Δ*t*_*window*_, *t*_*curr*_] is drawn around the current timestamp. The average for each bin (more exactly, position bin) in the distribution is taken across the window. If representation in the target region exceeds a threshold and representation in the off-target region is below a (different) threshold, a remote representation is considered to have occurred. Note that in this implementation we did not use the continuity constraint that combines information from the previous timestep with the spiking information from the current timestep. We also did not include a requirement for a specific LFP signature, although there would likely be some overlap among the events detected by this method and those that detect SWRs or population burst events (PBEs). Some example events are shown in Fig. 4b.

One pilot animal performed this task and demonstrated that the closed-loop neurofeedback worked as designed to detect and reward sustained remote representations (Fig. 4b). After several days of preliminary neurofeedback, we introduced the requirement that at least 2 tetrodes had spatially specific activity for event detection, so as to reward population-level representations. With this requirement, we found that detected event count increased across 9 feedback sessions targeting arm 1 and then after switching the target region to arm 2, also increased across 9 more sessions (Fig. 4c). This provides a preliminary suggestion that remote hippocampal content can be enriched via reward-based neurofeedback. In subsequent experiments, we tested whether this finding was reproducible across multiple animals and robust to additional controls.

## Discussion

Decoding neural activity allows experimenters to assess the structure of ongoing neural representations. Here we present a real-time system that implements a clusterless state space decoder. The specific implementation was designed to decode spatial representations from the spiking of hippocampal neurons, but the system would be relatively easy to modify to decode other variables. We improved upon previous real-time clusterless decoders by writing the software in pure Python to ease the experience of users intending to extend and customize the system for their needs. Despite the use of Python, our system achieved comparable performance to previous systems written with a compiled language, demonstrating the viability of using an interpreted language for this specific application. We also demonstrate the system in a pilot experiment, illustrating how it can be used to detect the expression of specific hippocampal representations of space and to trigger experimental stimuli based on that expression.

One area for improvement is the scalability of our decoder. For best results, one hardware thread is needed for every electrode group. However, the channel counts of neural recording devices are continuing to increase, and even with systems that have many computing cores, the number of channels per device will exceed current CPU capabilities. One potential solution would be to leverage computing on GPUs so that a single hardware thread can support multiple electrode groups. In order to fully take advantage of the GPU, this strategy would likely require some non-trivial programming to convert the synchronous computing into asynchronous. Alternatively, it may be possible to develop dimensionality reduction approaches that make it possible to carry out real-time clusterless decoding from very high channel count probes, something that is currently only possible offline (Y. Zhang et al. 2024).

While these improvements can be made, the current `RealtimeDecoder` has numerous applications within the neuroscience sub-field of hippocampal physiology by assisting researchers to run many kinds of closed-loop experiments. Applications to other neuroscience sub-fields are also possible. For example one may wish to design a brain-computer interface which can decode memories its human operator is attempting to retrieve, and to subsequently react in real-time based on that decoding. The computational approach illustrated by `RealtimeDecoder` has the potential to inform solutions to other decoding-based problems in general. Overall we believe our work serves as a valuable tool to accelerate neuroscience research.

## Appendix

### Customization

Although we have endeavored to write a generalized software system for real-time decoding, it is difficult to anticipate every use case. In some scenarios customization may be necessary to fulfill the particular needs of the experiment. Here we cover different layers of customization and explain the source code at a deeper level so that the user can better understand the modifications that must be made for their specific application.

### Customizing the data acquisition interface

As explained previously our software can be likened to a client in a client-server architecture, where `RealtimeDecoder` is the client and the data acquisition system is the server which sends preprocessed data for decoding. The decoder will work out of the box with Trodes software; for other systems, the user will need to develop a `DataSourceReceiver` subclass to interface with their data acquisition. The data acquisition is not constrained to be one that streams live data, as in the case of Trodes. One may wish to feed pre-recorded data into `RealtimeDecoder`, by reading from file. The parent class is defined in Listing 3, and an explanation of the intended uses of each method follows.

**Listing 3.**
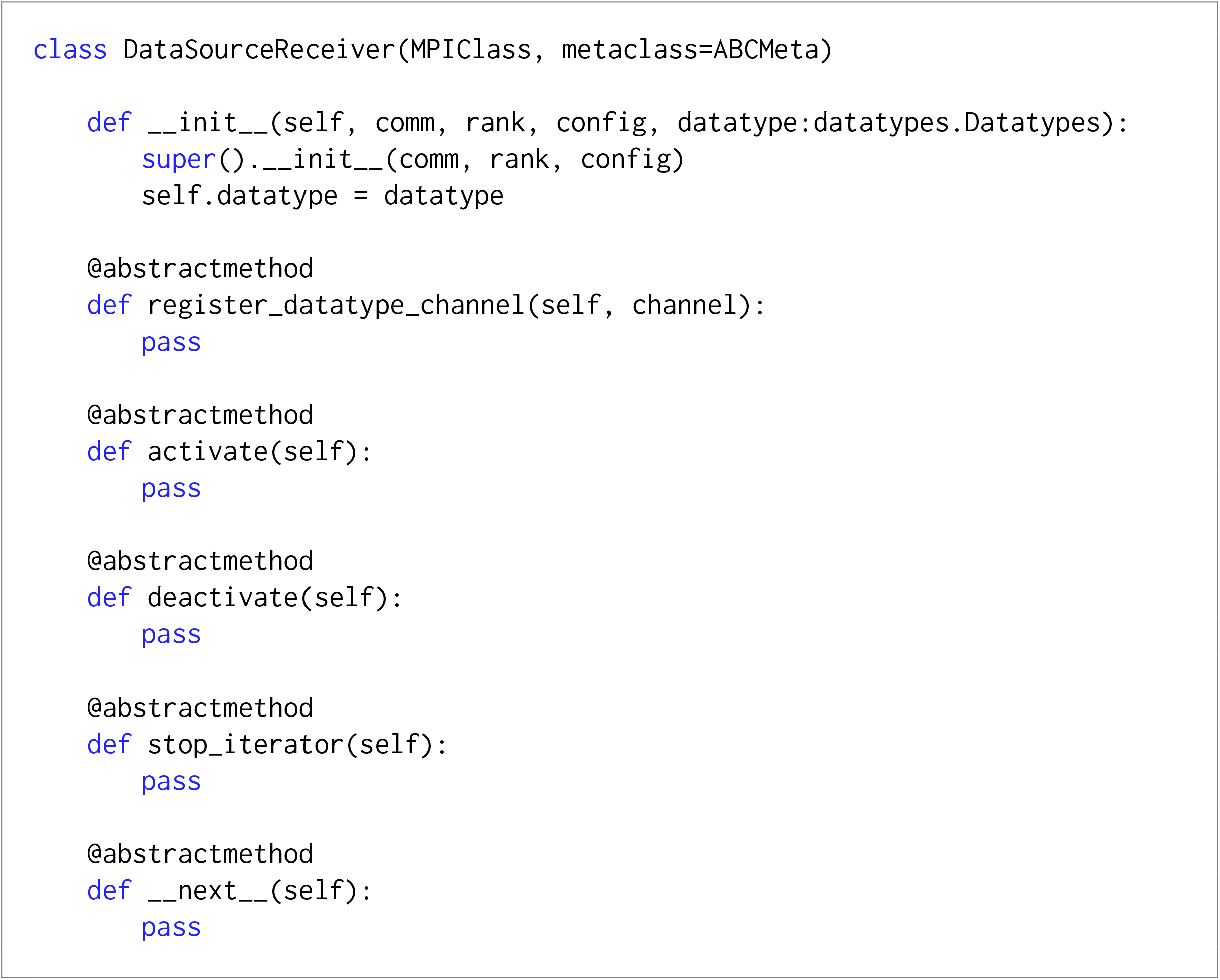
Definition of DataSourceReceiver class

In the `__init__()` method `comm` is the MPI communicator, `rank` is the MPI rank, and `config` is the dictionary obtained from parsing the configuration file (explained in the previous chapter). Finally `datatype` is one of the datatypes enumerated in the `datatypes` module.

The `register_datatype_channel()` method is used to add an electrode group that the object should receive from. Each electrode group may consist of multiple electrodes; for example a tetrode consists of 4 electrodes.

The `activate()` method signifies that the `DataSourceReceiver` is ready to receive data; thus `__next__()` may return a non-`None` object.

The `deactivate()` method signifies that the object is no longer receiving data. In this state `__next__()` should always return `None`.

The `stop_iterator()` stops the object from running entirely. Since the object can be used as an iterator (hence the `__next__()` method), this method should at the minimum contain a `raise StopIteration` statement.

Lastly `__next__()` should be called whenever data is to be returned. If no data are currently available or the object is in a deactivated state, then this method should return `None`.

Upon a call to `__next__()`, any `DataSourceReceiver` object that returns a data point in the following formats are automatically compatible with `RealtimeDecoder`. These are further explained below.

Spike data points are implemented as Listing 4. `timestamp` is a value that marks the timestamp of the data point and can be represented as a 32-bit unsigned integer. `elec_grp_id` is an integer denoting the electrode group the data point is coming from. `data` is the actual spike waveform data, a 2D array of shape (nch, nt) where nch is the number of electrodes in the electrode group and nt is the number of time samples in the spike waveform. `t_send_data` is a 64-bit integer marking the time in nanoseconds where the data was sent out by the data acquisition system. Similarly `t_recv_data` is a 64-bit integer marking the time in nanoseconds where the data was actually received by the `DataSourceReceiver`. Both `t_send_data` and `t_recv_data` are primarily used for diagnostics purposes to characterize network latency.

**Listing 4.**
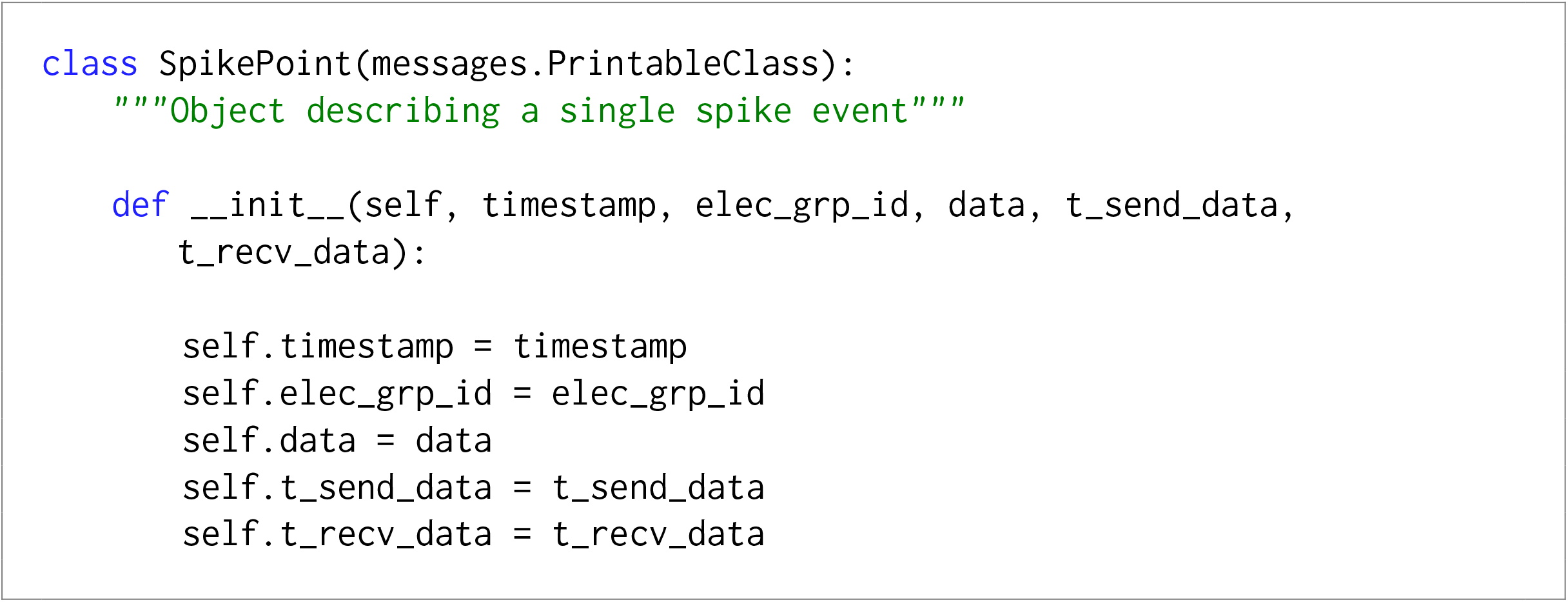
Definition of SpikePoint class

LFP data points are implemented as 5. For an `LFPPoint timestamp`, `t_send_data`, and `t_recv_data` have identical meanings as for a `SpikePoint`. Differences include `elec_grp_ids`: since a LFPPoint typically may contain data from *multiple* electrode groups `elec_grp_ids` is a list of those groups. Finally data is a 1D array containing the actual LFP data.

Lastly position data points are implemented as 6. Both `timestamp` and `t_recv_data` have identical meanings to those in `SpikePoint` and `LFPPoint`. In a real experiment the position is often coming from a head-tracked animal, so we will refer to that when explaining the rest of the data that this object represents. segment refers to a line segment (integer-valued) that the animal is closest to. `position` is a value in [0, 1] that denotes the position along the line segment. 0 and 1 are the ends of the segment. `x` and `y` are the x and y position of the animal, respectively, in pixels. `x2` and `y2` have the same interpretation as `x` and `y`, except that this is another point on the animal that is being tracked. For example the coordinates (x, y) and (x2, y2) may be the position of a red and green LED.

**Listing 5.**
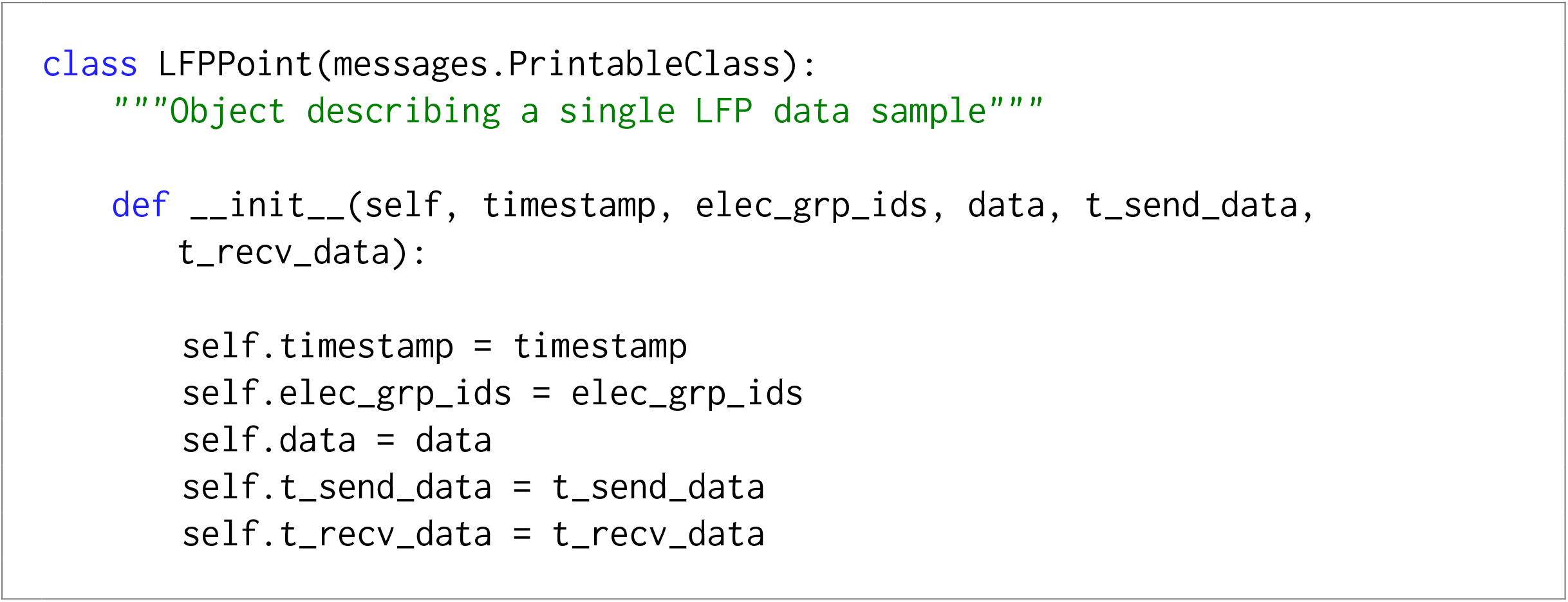
Definition of LFPPoint class

**Listing 6.**
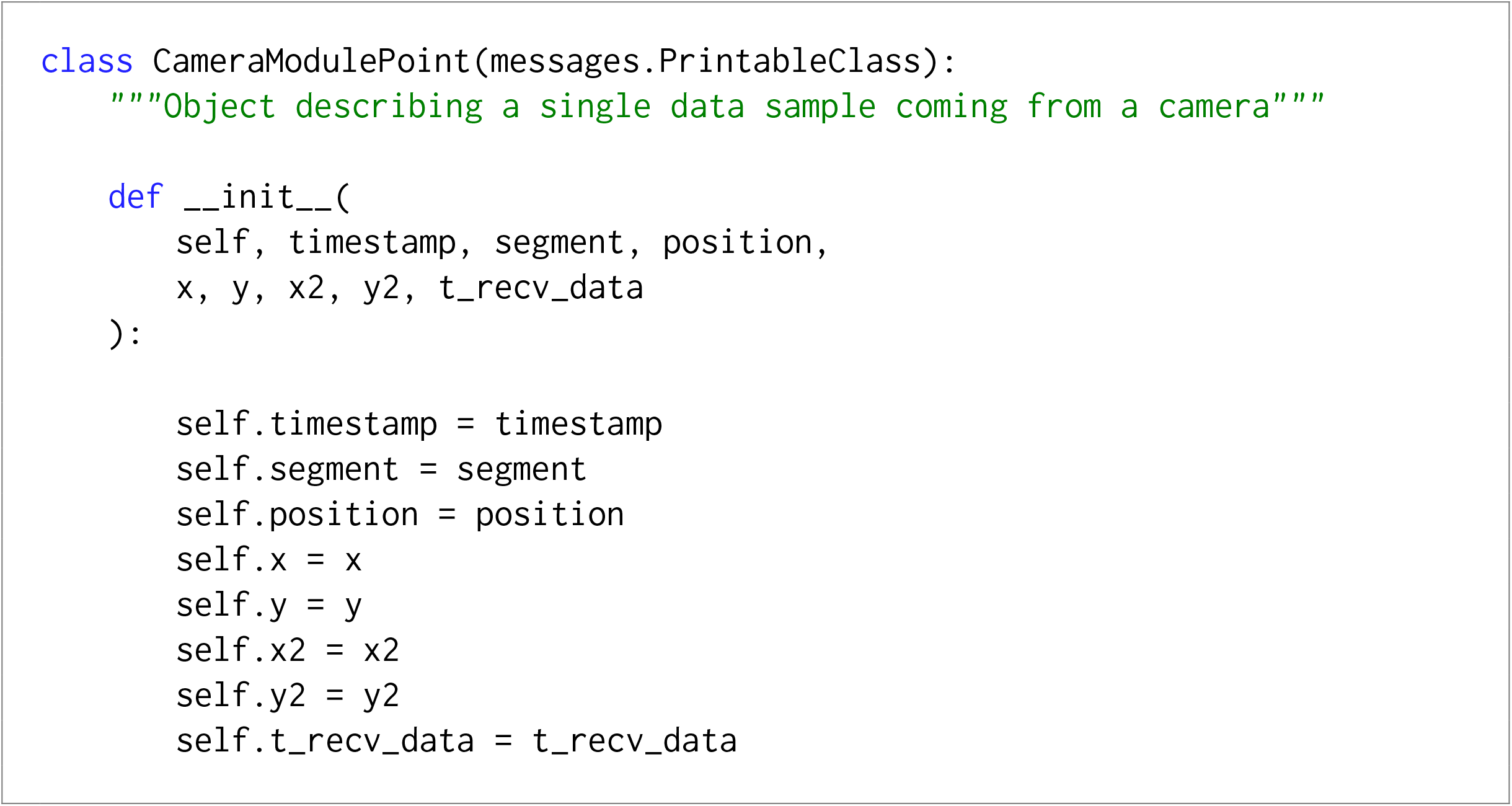
Definition of CameraModulePoint class

The fastest way to make `RealtimeDecoder` compatible with a custom data acquisition system is to develop a `DataSourceReceiver` subclass that still returns a `SpikePoint`, `LFPPoint`, or `CameraModulePoint`. If the experimenter desires to use a different data format, additional development will be necessary so that the software can handle the custom data objects. The most relevant modifications to make will be the `next_iter()` methods for the `RippleManager`, `EncoderManager`, and `DecoderManager` objects in ripple_process.py, encoder_process.py, and decoder_process.py, respectively.

### Customizing the record format

When `RealtimeDecoder` is running, it records results to disk. These can simply be thought of as log entries in binary form. For example a record is written when the posterior distribution is updated one time step.

Occasionally a user may wish to add, remove, or otherwise change the type of records that are written. At the minimum the appropriate `*Manager` object should be modified in its constructor. Listing 7 shows an example for the `DecoderManager`.

`rec_ids` are a list of integer-valued numbers, each of which describes the type of record that is being written. These are particularly useful when the outputs of `RealtimeDecoder` are merged into a single file. Different processes can write the same type of records, so when merging records, the ID is used to group them together.

`rec_labels` are used to label each element in a single record (i.e. a binary blob). This is useful for converting the binary data into human-readable format, such as a `pandas` dataframe.

Finally `rec_formats` describe the datatype (such integer, float, etc.) used to represent a given record. These are all format strings from the struct module in the Python standard library.

If a record is changed, the corresponding call to `write_record()` must likewise be updated so that the correct arguments are supplied. Other modifications to the code may be necessary. Since it is impossible to anticipate every possibility, such changes are not described here. Nevertheless this section should point the user in the right direction on where to begin their modifications.

**Listing 7.**
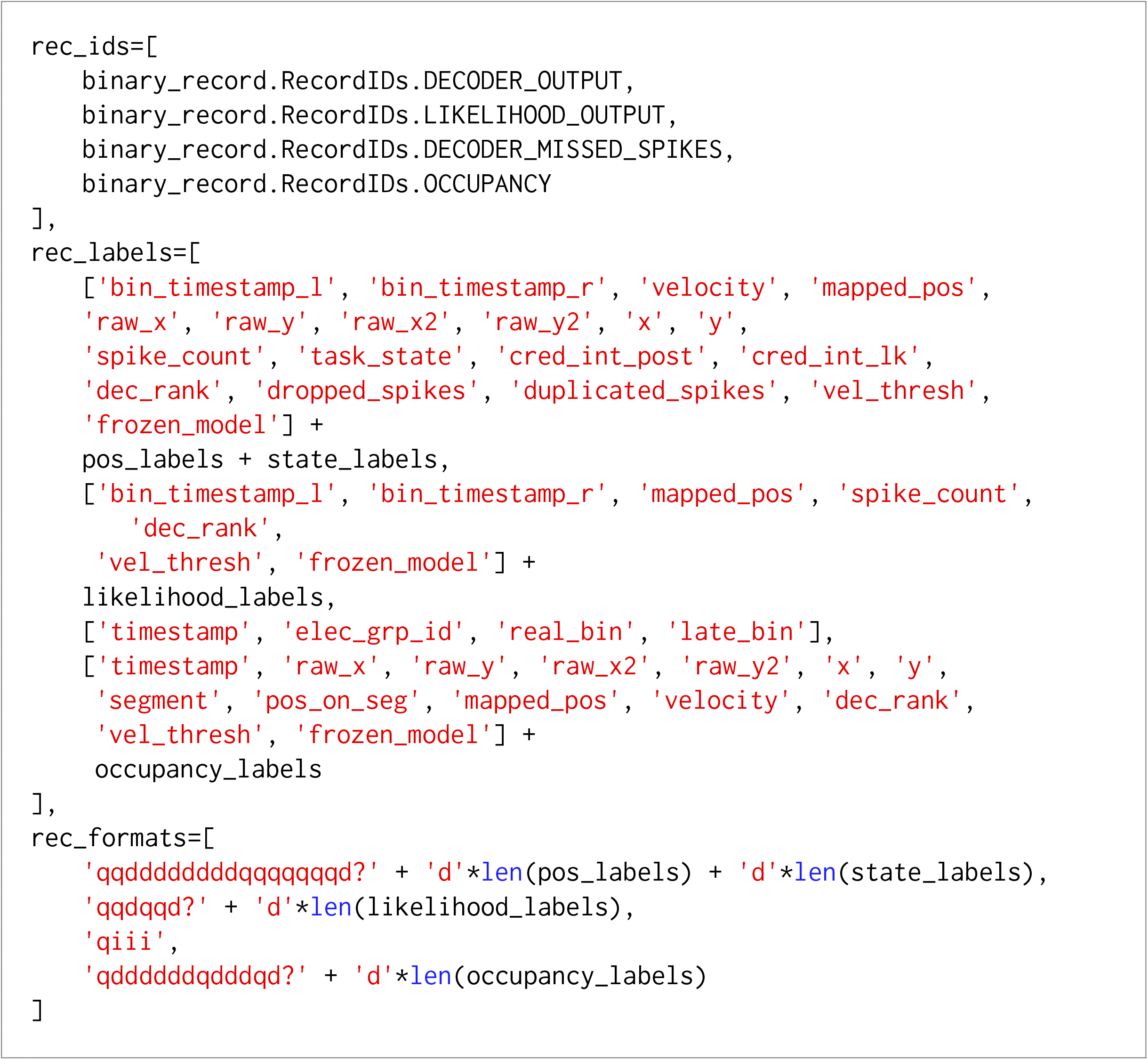
Defining record types

